# The Neural Correlates of Continuous Feedback Processing

**DOI:** 10.1101/2022.10.06.511117

**Authors:** Cameron D. Hassall, Yan Yan, Laurence T. Hunt

## Abstract

Feedback processing is commonly studied by analyzing the brain’s response to discrete rather than continuous events. Such studies have led to the hypothesis that rapid phasic midbrain dopaminergic activity tracks reward prediction errors (RPEs), the effects of which are measurable at the scalp via electroencephalography (EEG). Although studies using continuous feedback are sparse, recent animal work suggests that moment-to-moment changes in reward are tracked by *slowly ramping* midbrain dopaminergic activity. Some have argued that these ramping signals index state values rather than RPEs. Our goal here was to develop an EEG measure of continuous feedback processing in humans, then test whether its behaviour could be accounted for by the RPE hypothesis. Participants completed a stimulus-response learning task in which a continuous reward cue gradually increased or decreased over time. A regression-based unmixing approach revealed EEG activity with a topography and timecourse consistent with the stimulus-preceding negativity (SPN), a scalp potential previously linked to reward anticipation and tonic dopamine release. Importantly, this reward-related activity depended on outcome expectancy: as predicted by the RPE hypothesis, activity for expected reward cues was reduced compared to unexpected reward cues. These results demonstrate the possibility of using human scalp-recorded potentials to track continuous feedback processing, and test candidate hypotheses of this activity.

## 1 Introduction

Our daily lives often require us to monitor continuous feedback, yet much of what we know about feedback processing in the brain relies on experiments with discrete events only. For example, an unexpected reward is known to elicit phasic or “bursting” dopamine activity in monkey midbrain, which transfers back in time to a reward-predicting stimulus (Schultz et al., 1997, 2017). This is a pattern of activity consistent with a temporal difference reward prediction error (RPE), a computational measure of surprise (Sutton & Barto, 2018). In humans, this activity may be measurable at the scalp via electroencephalography (EEG) using the method of event-related potentials (ERPs). In particular, Holroyd and Coles (2002) proposed that reward-related changes in phasic midbrain dopamine modulate the activity of neurons in anterior cingulate cortex, resulting in an ERP component called the reward positivity (RewP^1^). The theory that RewP amplitude correlates with RPEs in humans has been influential, generating many experiments over the past 20 years (Glazer et al., 2018; Krigolson, 2017; Proudfit, 2015; Walsh & Anderson, 2012). However, the RewP is elicited by discrete feedback, so it is unclear whether such a signal would be present during continuous feedback processing.

In general, less is known about the neural correlates of continuous feedback processing compared to discrete feedback processing. Recent animal work has suggested that continuous feedback is tracked by tonic (“ramping”) dopamine as opposed to phasic dopamine. Specifically, a moving bar indicating the magnitude of an upcoming reward elicits tonic dopamine activity in the monkey midbrain (Wang et al., 2021). This is a significant result because it means that dopamine flexibly tracks both discrete and continuous feedback, and is consistent with other reports arguing that ramping dopamine activity can be described as a change in *state value* over time, tied to motivation (Hamid et al., 2016; Mohebi et al., 2019). *State* here refers to important information about the environment, such as a maze location or a cue, and *state value* is the future expected reward associated with a particular state. An RPE occurs when the state value is better or worse than expected. Some have argued that state values and RPEs map onto different types of dopamine activity (tonic and phasic, respectively: Hamid et al., 2016; Mohebi et al., 2019). However, others have argued that tonic and phasic dopamine are *both* responsible for signalling RPEs (Kim et al., 2020; Mikhael et al., 2022).

Although tonic dopamine may be a suitable candidate mechanism for tracking continuous feedback, it is currently unclear whether such activity could be measured using EEG. However, one reason to be hopeful comes from reward anticipation studies. Reward anticipation and consummation are traditionally dissociated, yet they share a similarity; both are known to elicit tonic dopaminergic activity in monkeys (Fiorillo et al., 2003; Wang et al., 2021). Relevant here, reward anticipation in humans is associated with an ERP component in humans called the stimulus-preceding negativity (SPN). The SPN is “slow”, ramping negatively from several hundred milliseconds before the onset of feedback^2^ and its amplitude depends on reward magnitude and predictability. The SPN is enhanced when feedback is associated with monetary reward (Kotani et al., 2003) and scales with the uncertainty of the outcome. Outcomes that are 50% likely are associated with greater ramping compared to outcomes that are 75% likely (Catena et al., 2012; Fuentemilla et al., 2013). Likewise, the SPN tends to be greatest early in learning when outcomes are less predictable (Morís et al., 2013). Finally, SPN amplitude is lower in individuals who have decreased tonic dopamine levels due to Parkinson’s disease (Mattox et al., 2006) or genetics (Foti & Hajcak, 2012).

Our goal here was to build on previous animal work (Kim et al., 2020; Wang et al., 2021) by identifying and testing a neural signature of continuous feedback processing in humans. In particular, we sought to capitalise upon recent developments in EEG analysis that use a regression-based approach to unmix components for continuously varying task variables from other discrete events (Ehinger & Dimigen, 2019; Hassall et al., 2022; Ruesseler et al., 2022; Crosse et al., 2016). We predicted that the resulting signal would resemble the SPN, an ERP component previously linked to reward anticipation and tonic dopamine (Glazer et al., 2018). Unlike previous work, we sought to understand how the brain tracks moment-to-moment changes in reward, not how it anticipates or processes a discrete reward. To do this, we designed a novel decision-making task in which the value of a reward cue changed continuously. We also varied outcome expectancy, which allowed us to test whether the identified signal tracks state values (Hamid et al., 2016; Mohebi et al., 2019) or RPEs (Kim et al., 2020; Mikhael et al., 2022). A signal that tracks state values (Hamid et al., 2016; Mohebi et al., 2019) ought to be greatest for continuous *predictable* reward cues because these cues become more strongly associated with the final outcome. Conversely, a signal that tracks RPEs ought to be greatest for continuous *unpredictable* reward cues because these cues would be more surprising overall.

## 2 Method

### 2.1 Participants

Twenty-one participants (5 male, 2 left-handed, *M_age_* = 25.81, *SD_age_* = 4.42) with normal or corrected-to-normal vision took part in the experiment. Participants provided informed consent approved by the Medical Sciences Interdivisional Research Ethics Committee at the University of Oxford. Following the experiment, participants were compensated £20 (£10 per hour of participation) plus a mean performance bonus of £2.02 (*SD* = £0.53). Data for one participant was excluded from all analyses due to the presence of many EEG artifacts (see Section 2.4.2 for details).

### 2.2 Apparatus and procedure

Participants were seated approximately 660 mm from a 599 mm X 337 mm display (60 Hz, 1920 by 1080 pixels, Acer XB270H, New Taipei City, Taiwan). Visual stimuli were presented using the Psychophysics Toolbox Extension (Brainard, 1997; Pelli, 1997) for MATLAB 2022a (Mathworks, Natick, USA). Participants were given written and verbal instruction to minimize head and eye movements as much as possible.

Participants played “Gnomes”, a continuous stimulus-response learning task. Participants were told that six gnomes, attending a fair, competed in a game of strength called “high striker” that involved striking an apparatus with a hammer to cause a puck to rise. The participants’ task was to bet on how high the puck would rise for each gnome (i.e., to bet on the strength of each gnome). Bets were placed by using a mouse to select a location on a vertical rectangular outline, representing the high striker apparatus. Participants then watched as a solid red rectangle, representing the location of the “puck”, gradually increased in height. Participants were awarded points depending on the how close their guess was to the actual outcome. Each gnome was encountered 25 times, in random order, for a total of 150 trials. See Supplementary Figures S1 and S2 for screenshots of the participant instructions.

Each trial was preceded by a centrally presented fixation cross subtending 1° of visual angle. After 400-600 ms (uniform distribution), one of six gnomes appeared for 1500 ms. The size of the gnomes varied in width and height; all dimensions were within 3.5°. The gnome was then reduced in size by half and moved to the lower portion of the display, signalling the participant to respond. Directly above the gnome, participants saw the centrally presented black outline of a vertical rectangle (1° wide and 3° high). After using the mouse to move a 2° horizontal black line up or down along the vertical rectangle, participants clicked the left mouse button to indicate the location of their guess. Following a 400-600 ms delay (uniformly distributed), participants observed the outcome of their bet. A red rectangle, 1° in width and sharing a lower border with the black rectangle, increased in height at a rate of 1° per second (like an increasing progress bar). The animation continued until the actual outcome was reached, then stopped. The participant could then see how close their guess was to the actual outcome (i.e., the difference between the horizontal bar and the final height of the red rectangle). This outcome remained on the display for 1000 ms (Figure 1a). Points in each trial were determined according to the difference between the participant’s guess and the actual outcome (in pixels), expressed as a proportion of the vertical black rectangle *p*, and converted to a value from 1–100 (more points for closer guesses) according to the formula 100*(1-p).Points were converted to a bonus payment at a rate of £0.0002 per point (*M_points_* = 10,088, *SD_points_* =2,670).

**Figure 1.**
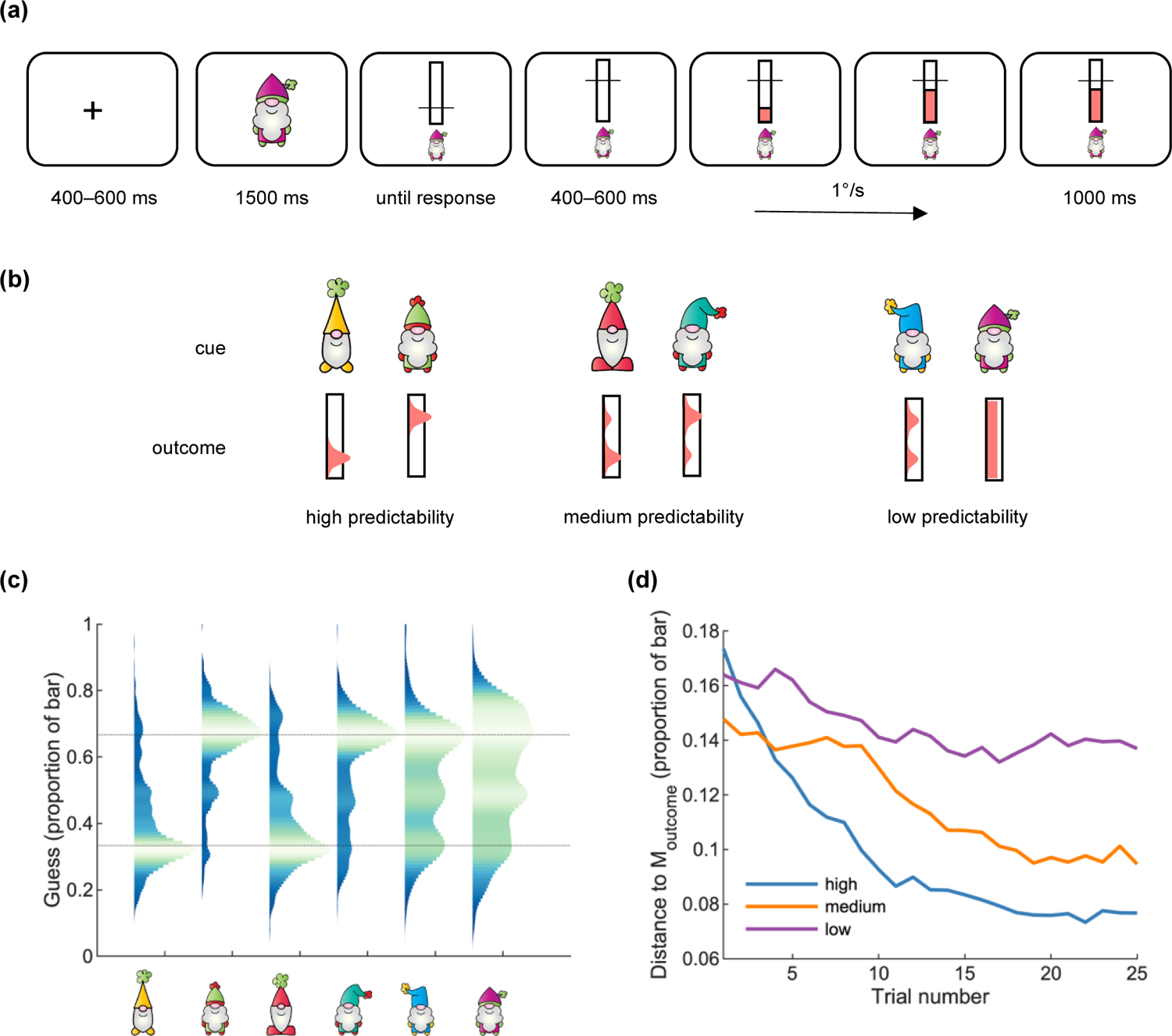
Participants learned to predict a continuous outcome. (a) Sample trial. Participants made a prediction about how high an animated bar would rise, then observed the outcome. (b) Outcome predictability according to a pre-trial cue. (c) Probability density estimates indicated that participants could predict the outcomes, as did (d) the distance between their guess and the mean outcome. The dotted lines in (c) indicate the mode of the “low” and “high” outcome distributions.

Each pre-trial cue (gnome) had a different outcome probability distribution, assigned randomly at the start of the experiment, meaning that some gnomes had a high level of predictability (and consequently, high reward expectancy), whereas others had medium or low levels of predictability (and medium/low reward expectancy). For two pre-trial cues (“high” predictability) outcomes were drawn from a Gaussian distribution (*SD* = 0.01) with either a low mean (1/3 of the vertical rectangle) or high mean (2/3 of the vertical rectangle). The same two distributions were combined for the “medium” predictability pre-trial cues using the MATLAB function *gmdistribution*: 80% low, 20% high for one pre-trial cue, and vice-versa for the other. Finally, there were two “low” predictability pre-trial cues, following either a 50-50 mixture of low and high Gaussians, or a uniform random distribution drawn from 0.2–0.8 of the vertical rectangle. See Figure 1b for a visual representation of the different outcome probabilities.

### 2.3 Data Collection

On each trial, the experimental software recorded the pre-trial cue identity, the participant guess (as a proportion of the height of the vertical black rectangle), and the outcome. As well, thirty-two channels of EEG, referenced to electrode Fz, were recorded using Brain Vision Recorder (Version 1.23.0003, Brain Products GmbH, Gilching, Germany. Thirty of the electrodes were place on a fitted cap (EASYCAP GmbH, Wörthsee, Germany) with a standard 10-20 layout. Two electrodes were also attached to the left and right mastoids. Conductive gel was applied to the electrodes to lower their impedance before recording. The EEG was sampled at 1000 Hz and amplified (actiCHamp Plus, Brain Products GmbH, Gilching, Germany) with a 280 Hz anti-aliasing filter.

### 2.4 Data Analysis

#### 2.4.1 Behavioural analysis

The behavioural data was analyzed in MATLAB 2022a (MathWorks, Natick, USA). To visualize participant guesses, we collapsed across all participant responses and used the *distributionPlot* function to estimate the guess probability density associated with each pre-trial cue (Jonas, 2008). For each participant, outcome predictability (high, medium, low), and trial (1-25), we computed the mean error, defined as the distance between the participant’s guess and the actual outcome (expressed as a proportion of the total bar). Next, we averaged across trials to get a single distance for each participant and outcome predictability. We then computed the continuous predicted reward experienced by each participant across the entire task, which was required for our EEG analysis. The time resolution of the continuous predicted reward was 250 Hz, matching that of the downsampled EEG. During non-reward times the continuous predicted reward was set to zero. Finally, to quantify reward expectancy for each participant and pre-trial cue we computed the average reward that was experienced on a trial-by-trial basis (i.e., for a given cue and trial we computed the average reward over that cue’s previous trials). The resulting signal was used as a parametric regressor in our final EEG analysis.

#### 2.4.2 EEG preprocessing

The EEG was also analyzed in MATLAB 2022a (MathWorks, Natick, USA) using the EEGLAB library (Delorme & Makeig, 2004). After downsampling to 250 Hz, we applied a bandpass filter (0.1-30 Hz, 50 Hz notch), and re-referenced to the average of the mastoid signals. Ocular artifacts were then identified and removed using independent component analysis (ICA). The ICA was trained on 3-second windows extending 0.2 s prior to 2.8 s after the onset of each pre-trial cue. Epochs with large artifacts (a range in potential of more than 500 mV) were excluded from the ICA. Ocular components were identified using the function *iclabel* and removed from the continuous dataset if assigned an “Eye” label with greater than 0.8 likelihood (Pion-Tonachini et al., 2019).

#### 2.4.3 Correlation analysis

Our initial approach was to simply compute the zero-lag correlation between the continuous reward signal (Figure 2a) and the continuous preprocessed EEG. We first identified artifacts in the continuous EEG using the *uf_continuousArtifactDetect* function from the Unfold toolbox (Ehinger & Dimigen, 2019), which is based on code from the ERPLAB toolbox (Lopez-Calderon & Luck, 2014). This sliding-window approach (window size: 2000 ms, step size: 100 ms) flagged all samples in a window with a peak-to-peak difference of more than 150 mV. Flagged samples were then removed from subsequent analysis. One participant had excessively noisy EEG and was excluded from all analyses (17.51% of modelled samples). Of the remaining 20 participants, we removed an average of 2.49% (*SD* = 2.19%) of modelled samples (that is, samples corresponding to non-zero elements of the continuous reward). A Pearson correlation coefficient (Pearson’s *r*) was then computed for each participant and electrode (Figure 2b).

**Figure 2.**
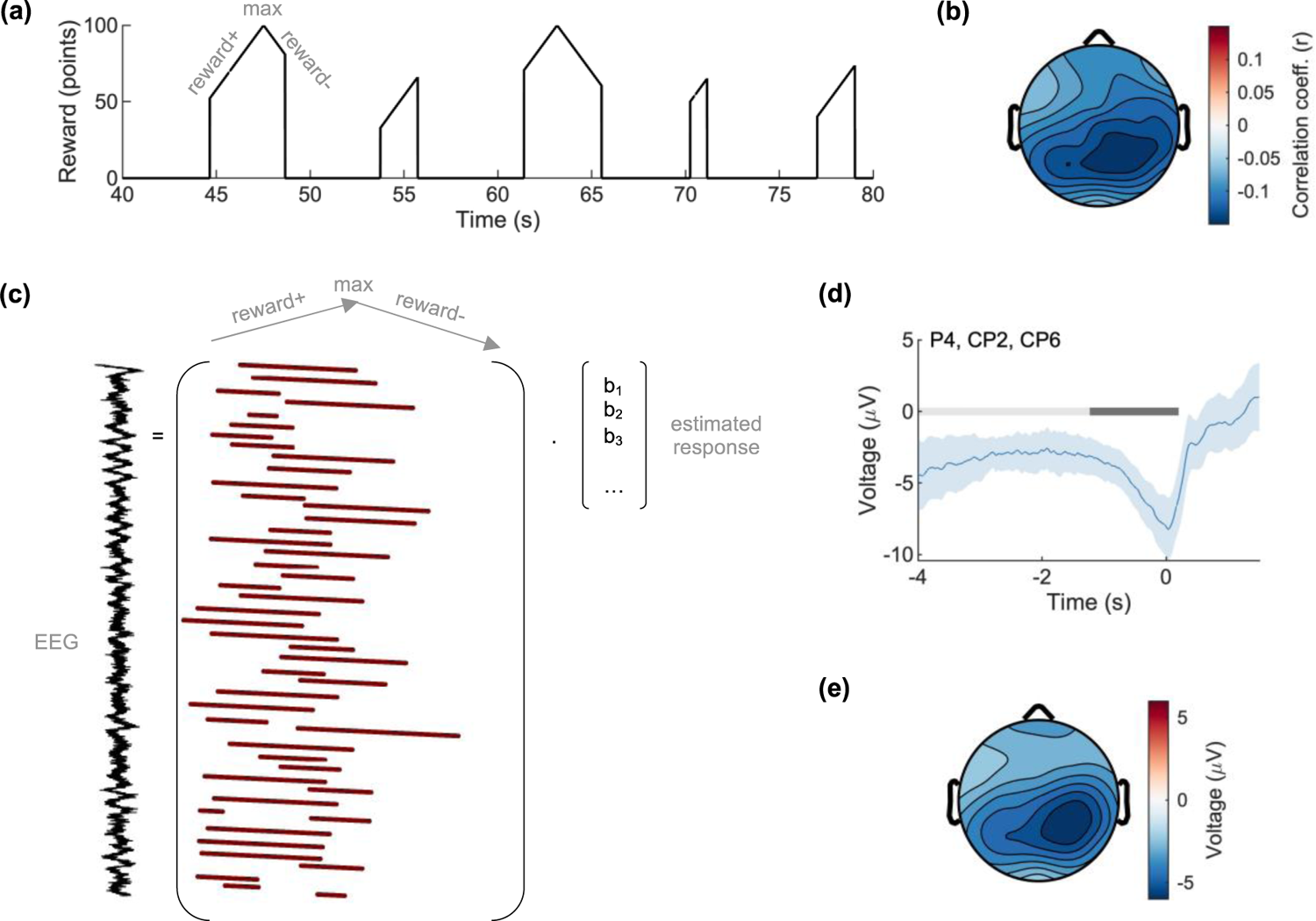
EEG tracks continuous predicted reward. Across the entire recording session, continuous reward (a) correlated maximally with EEG at a central-parietal scalp location (b). The EEG at rewarded timepoints was then modelled using linear regression. Regressors (1 during modelled times and 0 otherwise) were shifted to align to the time of maximum possible reward. Beta values *b* corresponded to the reward-related response which, when estimated, revealed a negative-going deflection (d) that was again central-parietally located (e). The shaded region indicates a 95% confidence interval, and the grey bars show the significant temporal clusters (light grey: without baseline correction, dark grey: with baseline correction).

#### 2.4.4 Regression analysis

The scalp topography in Figure 2b is useful because it indicates a relationship between scalp EEG data and continuous reward prediction. However, there are two issues with this analysis. First, it only examines the relationship between EEG and continuous reward at a single time lag (zero-lag correlation). In other words, a correlation analysis is uninformative about how the neural response to continuous reward prediction evolves over time. Second, a correlation analysis does not correct for other components that are also likely present in the EEG, such as movement-related activity at the onset and offset of the animated bar. To better quantify how EEG tracks continuous reward, we therefore constructed a *reward-aligned* ERP for each participant. To account for possible component overlap, we used a regression-based approach in which the ongoing EEG is modelled as a linear sum of underlying “regression-ERPs” or rERPs (Burns et al., 2013; Ehinger & Dimigen, 2019; Smith & Kutas, 2015a, 2015b). By “reward-aligned”, we mean that the rERP was time-locked to the time of maximum possible reward (that is, the time at which the rising bar matched the participant’s guess for “too low” guesses or *would have* matched the participant’s guess for “too high” guesses). This ensured that regressors represented the same instantaneous reward value across all trials. In practice, we did this by shifting the reward-related regressors earlier or later in time (see Figure 2c). The amount of shift varied from trial to trial, depending on the participant’s guess.

In our first regression analysis, we collapsed across pre-trial cue types and included regressors corresponding to three “events”: 0–800 ms relative to the start of the bar animation, the continuous reward signal described above, and 0–800 ms relative to the end of the bar animation when the participant learned the final outcome. Note that the design matrix in Figure 2c only shows the reward-related regressors. The purpose of this analysis was to verify the presence of a continuous reward signal. No baseline correction was applied initially. However, following visual inspection of the resulting signal, a −4000 ms to −1000 ms baseline was subtracted from the beta estimates to better isolate the prominent negativity observed around the time of maximum possible reward (i.e., the mean voltage in this time range was computed for and subtracted from the rERP for each participant and electrode).

Next, to determine the effect of outcome expectancy, we modelled the same three events (animation start, continuous reward, animation end) separately for each outcome predictability (high, medium, low). Finally, as outcome predictability was learnt gradually across the block, we sought to better measure (and isolate) the effect of outcome expectancy by collapsing across pre-trial cue types as before but including the trial-by-trial average reward for each pre-trial cue as a parametric regressor (see Section 2.4.1).

We used regularization to reduce model overfitting. Overfitting of noise was a concern because the number of modelled timepoints varied throughout the epoch, e.g., there were fewer timepoints towards the end (See Supplementary Figure S3). We used a first-derivative form of Tikhonov regularization (Kristensen et al., 2017), which imposes a smoothness constraint (Reichel & Ye, 2008). Ten-fold cross-validation was used to select an optimal regularization parameter for each participant. The error for a particular participant and fold was defined as the average mean-squared error across all electrodes. The following lambda values were tested: 100, 1000, 10000, 100000, 1000000. The optimal lambda for each participant minimized the mean fold error across all ten folds (see Supplementary Figure 4).

#### 2.4.5 Statistics

To check whether participants learned to predict trial outcomes we compared the mean distance between response and outcome via one-way repeated-measures ANOVA. Partial and generalized eta-squared were computed as:

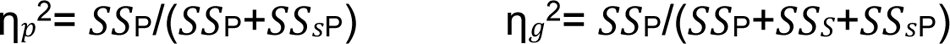

where *SS*_P_ was the sum of squares of the predictability effect (high, medium, low), *SS*_sP_ was the error sum of squares of the predictability effect, and *SS*_S_ was the sum of squares between subjects.

To check whether continuous EEG correlated with continuous reward, we conducted a repeated-measures *t*-test of the correlation coefficient at each electrode site. A Bonferroni-corrected alpha value of .05/30 = .0017 was used. Effect sizes were computed according to:

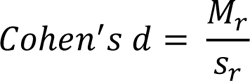

where *M_r_* was the mean of the correlation coefficients and *s_r_* was the standard deviation of the correlation coefficients (Cumming, 2014).

For our regression analyses, we used cluster-based permutation testing (Maris & Oostenveld, 2007). For each electrode, we computed a single-sample *t*-statistic at each sample point to determine whether the voltage at that sample point differed from zero. We then identified clusters of sample points for which the *t*-values exceeded the 2.5th or 97.5th percentile. Spatial clusters were defined according to an electrode template from the FieldTrip toolbox (Oostenveld et al., 2010). For each spatial cluster, we defined a “cluster mass” as the sum of the absolute values of the temporal cluster *t*-values; to be included in a cluster mass, the voltage at a sample point had to reach significance for all members of the spatial cluster. We restricted the search for temporal clusters to the window −4000 ms to 200 ms relative to the time of maximum possible reward. As Supplementary Figure S4 shows, most modelled sample points occurred prior to the time of maximum possible reward (i.e., the animated bar did not actually reach the participant’s guess). The interval −4000 to 200 ms ensured that there were at ten least modelled sample points per condition, on average. To determine whether the observed cluster masses exceeded what could occur by chance, we permuted the participant rERPs by randomly flipping the entire signal at all electrodes in the vertical axis. This is equivalent to swapping condition labels if one condition is “signal” and one condition is “baseline” (where the “baseline” is a vector of zeros). We then computed the cluster masses of the permuted waveforms and recorded the maximum cluster mass (or zero if there were no clusters). We tested 1000 permutations in total. Finally, we labelled an observed cluster as “significant” if its cluster mass exceeded 95% of the permuted cluster masses. The reported *p*-value was the proportion of permuted cluster masses exceeding the observed cluster mass. For each significant cluster, we also reported an effect size by averaging the EEG over the cluster electrodes and sample points for each participant and computing:

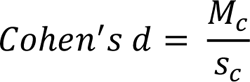

where *M_c_* and *s_c_* were the mean and standard deviation of the resulting cluster voltages.

## 3 Results

### 3.1 Behavioural Results

To identify the neural correlates of continuous feedback processing, we designed a novel task in which participants tried to predict the outcome of an animated rising bar (the final position). After indicating their guess, participants watched the bar rise and were awarded a point amount that depended on the distance between their guess and the outcome. Outcome predictability varied depending on a pre-trial cue that appeared prior to each round (Figures 1a and 1b). After pooling responses from all participants, the estimated guess distribution suggested that participants were able to match their guesses to the mean outcome (Figure 1c). Examining the mean distance between guesses and outcomes revealed an effect of outcome predictability (high, medium, low), *F*(2,38) = 13.09, *p* < .001, η_p_ ^2^ = 0.41, η_g_ ^2^ = 0.18. See Supplementary Table S1 for the mean distances.

### 3.2 EEG Results

#### 3.2.1 Correlation results

We first tested whether and how the time-varying continuous prediction of reward correlated with the time-varying EEG signal at zero lag. There was a negative correlation between continuous EEG and continuous reward at most electrode sites (Figure 2b). The correlation coefficients peaked at electrode CP2, Pearson’s *r* = −0.14, 95% CI [−0.18, −0.11], where it differed significantly from zero, *t*(19) = −8.24, *p* < .001/30 (Bonferroni-corrected for 30 comparisons).

#### 3.2.2 Regression results

Next, we modelled event-related activity relative to the maximum possible reward (i.e., when the animated bar crossed or would have crossed the participant’s guess) – see Figure 2c. This revealed a negative-going reward-related signal at a cluster of electrodes centred on Pz (Pz, CP1, CPz, CP2) spanning −4000 ms to 200 ms relative to the time of maximum possible reward (the entire tested interval), *p* < .001, Cohen’s *d* = −1.68. The effect remained after baseline correction but was shifted rightward on the scalp to a cluster centred on electrode P4 (P4, CP2, CP6) and limited to −492 ms to 192 ms (*p* < .001, Cohen’s *d* = −0.66) – see Figures 2d and 2e. The neural responses to the start and end of animation were not analyzed further but can be seen in Supplementary Figure S5.

We then tested whether this time-varying reward signal might be modulated by outcome predictability, as manipulated by the different pre-trial cues. After splitting by outcome predictability, we observed a difference (high versus low predictability) at a cluster located at Pz (Pz, CP1, CPz, CP2) from −516 ms to −164 ms relative to the time of maximum possible reward, *p* = .007, Cohen’s *d* = −0.70 (Figure 3). Consistent with an RPE account of this signal, we found it was smallest when the outcome was most predictable (high predictability condition).

**Figure 3.**
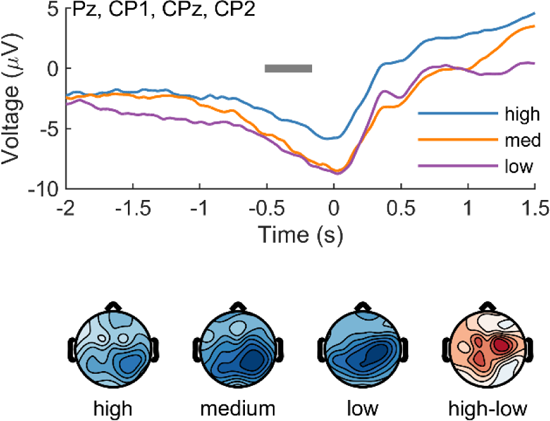
Continuous reward processing depends on outcome predictability. When outcome predictability was high, the preceding reward-related activity was reduced relative to the less predictable outcomes (medium, low). The grey bar indicates the significant temporal cluster (high versus low).

Finally, we measured the effect of outcome predictability in a different way: by examining the parametric effect of reward *expectancy*. We included an “expectancy” parametric regressor, quantified as the average trial-to-trial reward associated with each pre-trial cue (see Section 2.4.1). We replicated the mean reward-related signal from our first GLM – a cluster centred on Pz (Pz, CP1, CPz, CP2) from −4000 ms to 200 ms, Cohen’s *d* = −1.54. Applying a −4000 ms to −1000 ms baseline as before yielded a significant cluster at the same location but from −468 ms to 192 ms, Cohen’s *d* = −0.92 (Figure 4a). As further evidence of an RPE signal, we observed a frontal signal related to reward expectancy at a cluster centred on F4 (F4, FC2, FC6) from −1692 to −544 ms relative to the time of maximum possible reward, *p* < .001, Cohen’s *d* = 0.88 (Figure 4b).

**Figure 4.**
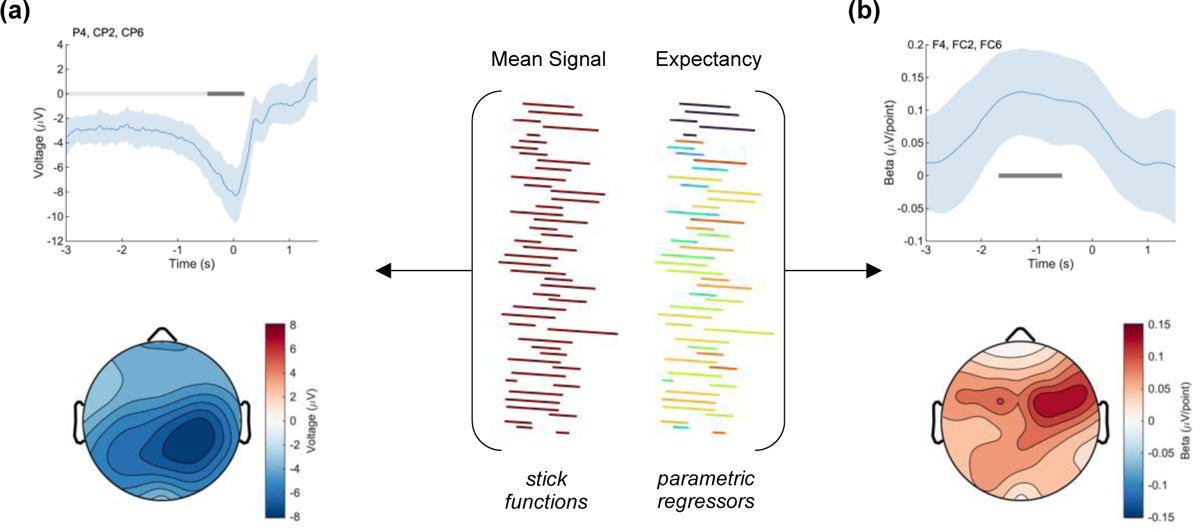
Parametric effect of expectancy on reward processing. We decomposed the reward signal into a parietal effect (a) plus a frontal parametric effect tracking expectancy, defined as the mean trial-to-trial reward associated with each cue (b).

## 4 Discussion

Our understanding of human feedback processing mostly relates to the anticipation and processing of discrete outcomes. However, recent animal work has suggested that a complete understanding of feedback processing should also include continuous outcomes (Kim et al., 2020; Mikhael et al., 2022; Wang et al., 2021). We built on this animal work by identifying a scalp-recorded signal in humans related to continuous feedback processing. In line with a previous study in rodents that monitored activity of dopamine neurons directly (Kim et al., 2020), the scalp EEG signal we found has properties consistent with a reward prediction error (RPE).

First, we observed that the amplitude of this continuous feedback signal varied inversely with expectancy – the more predictable the outcome, the smaller the signal (Figure 3). This result mirrors what has been found in discrete feedback studies (Holroyd & Krigolson, 2007; Sambrook & Goslin, 2015; Williams et al., 2017). Those studies found that the RewP (discussed earlier) is greatest for unexpected outcomes. This observation led to the view that the RewP tracks discrete RPEs via the effect of phasic dopamine on the anterior cingulate (Holroyd & Coles, 2002; Walsh & Anderson, 2012). Here we extend this view to include continuous feedback and speculate that both types of scalp-recorded signals reflect the activity of a common midbrain dopaminergic system – phasic activity for discrete outcomes, tonic activity for continuous outcomes. In line with this interpretation, we observed an effect of outcome expectancy at a frontal location consistent with the RewP (Figure 4). Taken together, our results support the claim that tonic dopamine tracks continuous temporal difference RPEs, not state values.

Further support for this claim is provided by Kim et al. (2020). There, mice moving through a virtual maze were unexpectedly “teleported” closer to their destination. Across a series of experiments, teleportation events elicited phasic midbrain dopaminergic activity consistent with temporal difference RPEs but inconsistent with state values. Regular progress towards the mouse’s goal was associated with tonic activity, as shown in previous work (Hamid et al., 2016; Mohebi et al., 2019; Wang et al., 2021). The authors reconcile these two types of activity by providing a computational account in which phasic and tonic dopaminergic activity share a common function: the signalling of moment-to-moment RPEs (Kim et al., 2020; Mikhael et al., 2022).

In the present study, participants were presented with a continuous reward cue that changed at a constant rate. In other words, there were no unexpected “jumps” analogous to the teleportation events in the Kim et al. (2020) study. Instead, we used pre-trial cues to vary outcome expectancy. This was done for methodological reasons. Unlike single-unit recordings, EEG activity reflects a mixture of underlying components (the *superposition* problem: Luck, 2014). We reasoned that a sudden jump in the level of the continuous reward cue would result in a visual evoked potential superimposed on reward-related activity. Such a mixed signal will need to be unmixed and quantified in order to fully test the predictions of Kim et al. (2020) in future EEG work.

The signal we identified also resembles the SPN, an ERP component previously associated with the anticipation of discrete outcomes. Both signals are slow, negative, and parietal. So, is the continuous signal that we identified just the SPN seen previously when presenting discrete, event-based feedback? Recall that each trial in our task concluded when the moving bar halted, and participants learned the outcome of the trial. Such an event could – if anticipated – have generated a discrete SPN independent of the continuous reward cue that we claim is the variable of interest here. In this case, participants could have ignored the moving bar altogether and simply focused on the relevant event for learning – the trial outcome. Note however that our trials lacked temporal predictability – knowing exactly *when* an event will occur. This differs from event uncertainty, discussed earlier (whether a reward will be received at all). Although the SPN is enhanced by event uncertainty (Catena et al., 2012), it is diminished by temporal uncertainty (Brunia et al., 2011). We suspect that our task lacks the temporal predictability required to elicit a discrete SPN, especially in the “low predictability” condition. Furthermore, we would expect any discrete SPN effects to be greatest in the “high predictability” condition – the opposite of what was observed (Figure 3).

For these two reasons – lack of temporal predictability and direction of effect – we are hesitant to identify our signal as merely a discrete SPN. However, the two signals may share a similar process that is revealed in two different ways. In the case of the discrete SPN, upcoming rewards are anticipated because they are temporally predictable. In the case of our “continuous SPN”, upcoming rewards do not occur after a set temporal interval but can nevertheless be tracked in real time via (in our case) an animated moving bar. The resulting signal shares a timing and topography with the SPN, as discussed. Another clue that these signals may share a similar underlying process comes from previous work on uncertainty (Catena et al., 2012; Fuentemilla et al., 2013). Consistent with our results, these studies each localized the effect of uncertainty on the SPN to frontal regions, either by examining the scalp (Fz: Fuentemilla et al., 2013) or through source analysis (ACC, DLPFC: Catena et al., 2012).

Real-world outcomes may be discrete or continuous, yet most experiments tend to focus on discrete outcomes only. This has resulted in an incomplete understanding of naturalistic feedback processing. We address this issue by identifying a neural signature of continuous feedback processing that, like its discrete counterpart, depends on outcome expectancy and is therefore consistent with a reward prediction error (RPE) signal. Our results complement a large body of work focused mainly on discrete outcomes and suggest a common dopaminergic mechanism underlying both discrete and continuous feedback processing.

## Author Note

This research was funded by a Natural Sciences and Engineering Research Council of Canada (NSERC) Postdoctoral Fellowship to Cameron D. Hassall (PDF 546078 - 2020), a Sir Henry Dale Fellowship from the Royal Society and Wellcome (208789/Z/17/Z) to Laurence T. Hunt., and a NARSAD Young Investigator Award from the Brain and Behavior Research Foundation to Laurence T. Hunt. This research was supported by the NIHR Oxford Health Biomedical Research Centre. The Wellcome Centre for Integrative Neuroimaging was supported by core funding from Wellcome Trust (203139/Z/16/Z).

For the purpose of Open Access, the author has applied a CC BY public copyright licence to any Author Accepted Manuscript version arising from this submission.

The authors have no conflict of interest.

## Author Contributions

**Cameron D. Hassall:** Conceptualization (equal); data curation; formal analysis; investigation (equal); methodology (equal); writing – original draft preparation; writing – review and editing (equal). **Yan Yan:** Investigation (equal); writing – review and editing (equal). **Laurence T. Hunt:** Conceptualization (equal); funding acquisition; methodology (equal); supervision; writing – review and editing (equal).

## Data Availability Statement

EEG dataset is available at https://doi.org/10.18112/openneuro.ds004262.v1.0.0. Analysis scripts are available at https://github.com/CCNHuntLab/continuousfeedback.

## Footnotes

1 Other names include the feedback-related negativity (FRN), the feedback error-related negativity (fERN), the medial-frontal negativity (MFN), and the feedback negativity (FN). See Proudfit (2015) for a detailed explanation. Also note that there has been debate about whether frontal cortex can be affected directly by dopamine or only indirectly via glutamate (Ullsperger et al., 2014).

2 In general, any anticipated stimulus ought to elicit an SPN. Furthermore, the SPN may be a subcomponent of another ERP component called the contingent negative variation (CNV). See van Boxtel and Böcker (2004) for a discussion.

