## Supplementary Material for "The Neural Correlates of Continuous Feedback Processing"

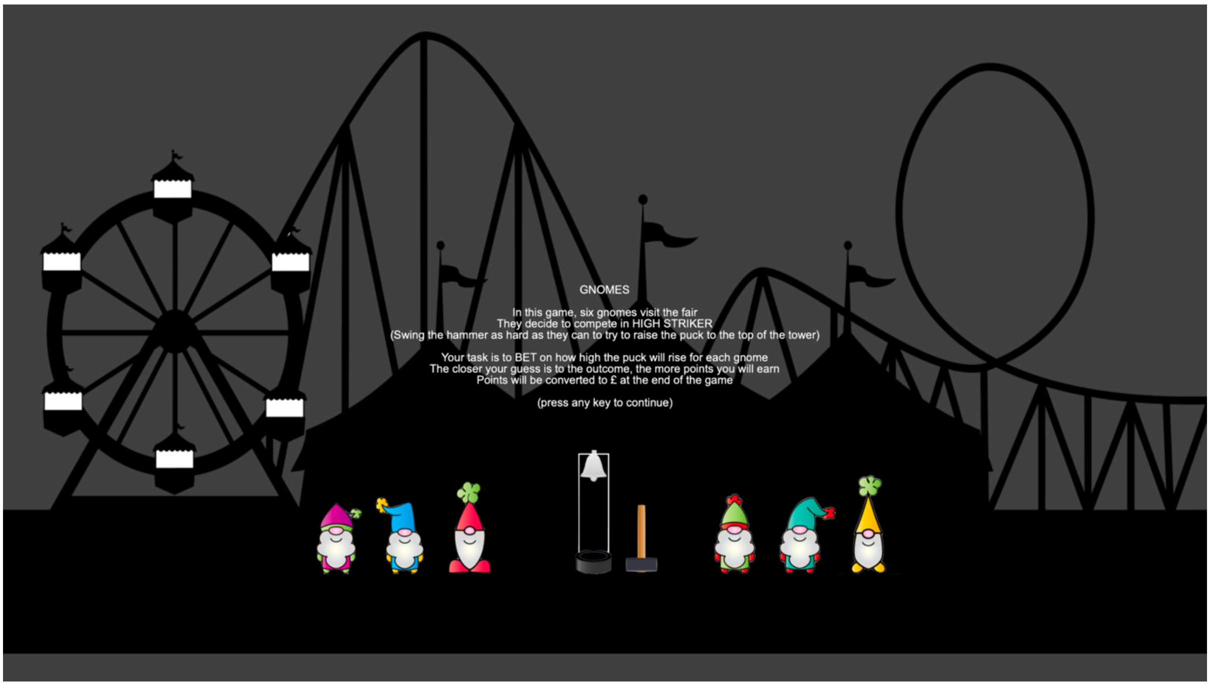


**Supplementary Figure S1.** First instruction screen.

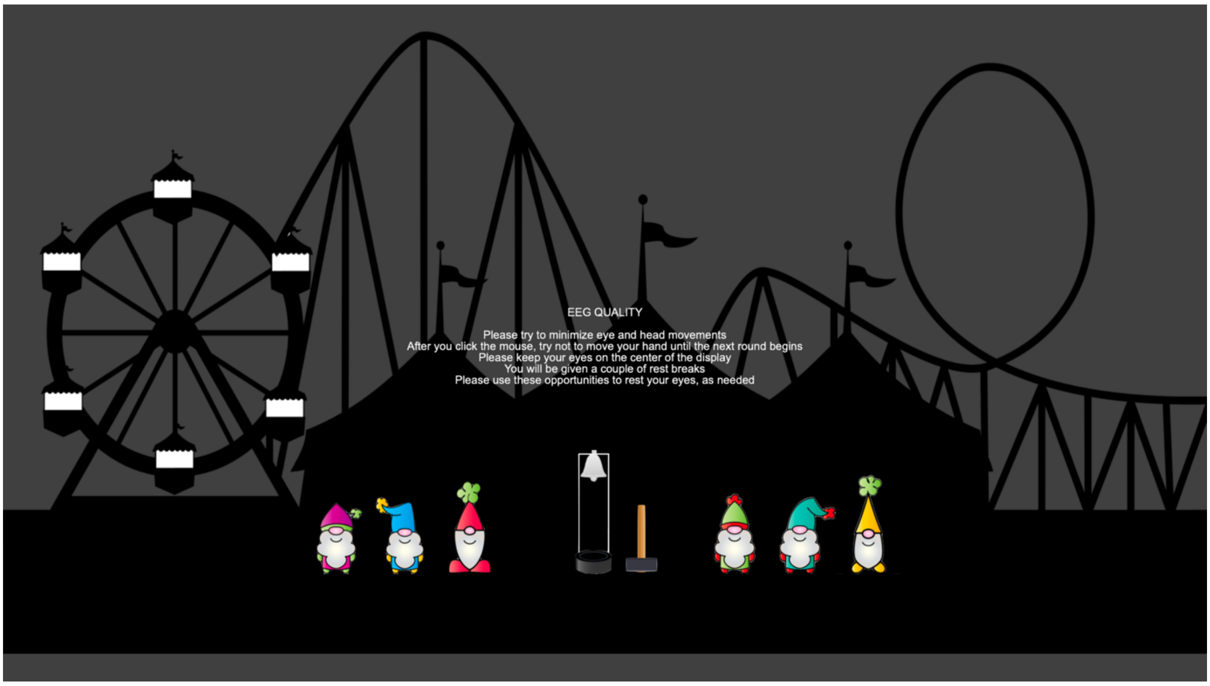


**Supplementary Figure S2.** Second instruction screen.

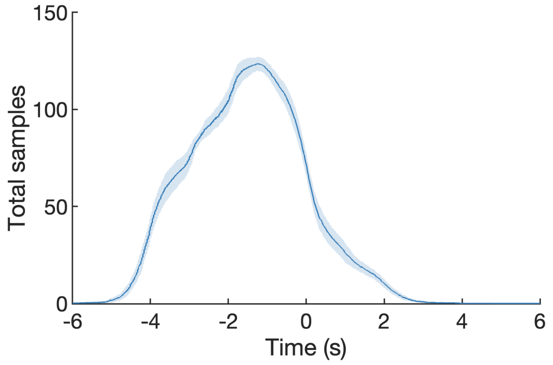


**Supplementary Figure S3.** Mean total number of modelled timepoints relative to the maximum reward value at 0 s. The shaded region indicates the 95% confidence interval.

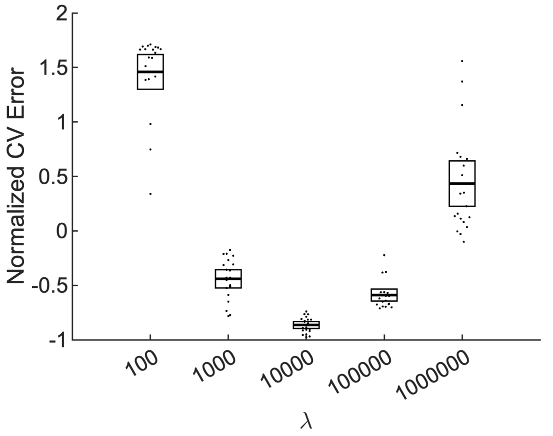


**Supplementary Figure S4.** Normalized cross-validation error. Dots represent individual participants, and error bars represent 95% confidence intervals. For all participants, the error was minimized by a lambda value of 10,000.

Supplementary Table 1

*Mean Distance Between Response and Outcome as a Proportion of Bar Height*

| Outcome Probability | Mean (Proportion) | 95% CI |
|---|---|---|
| High | 0.10 | [0.07, 0.12] |
| Medium | 0.12 | [0.10, 0.13] |
| Low | 0.15 | [0.13, 0.16] |

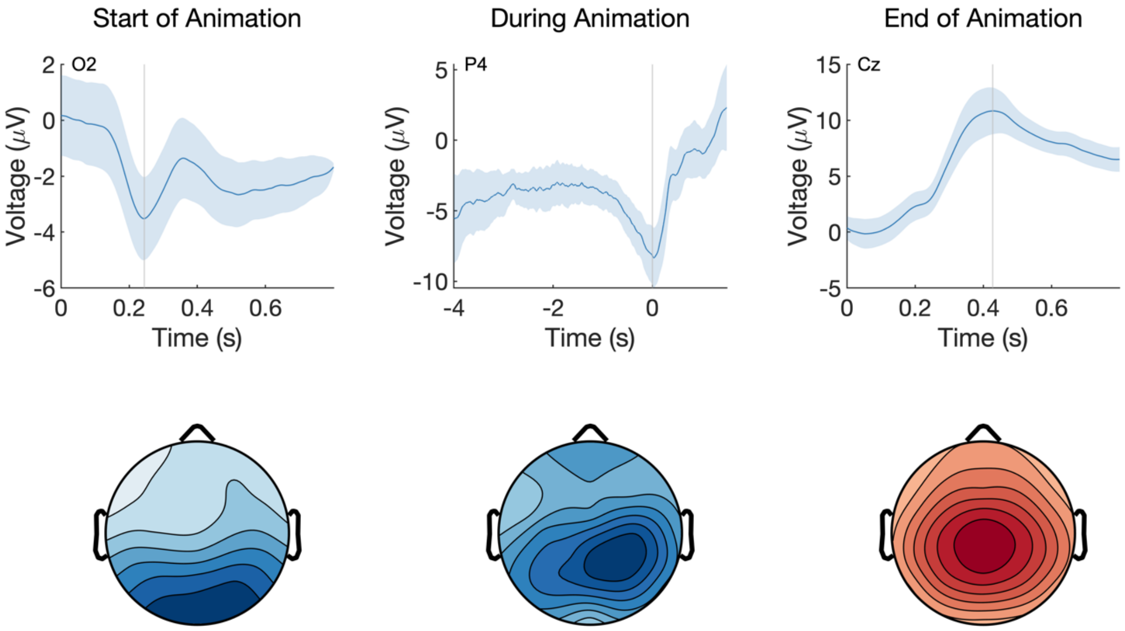


**Supplementary Figure S5.** ERPs before, during, and after the animation of the bar indicating the outcome of a trial. The plotted electrodes show the maximum signal location. The shaded regions indicate 95% confidence intervals. The scalp topographies represent the voltage at each electrode at the sample indicated by the vertical grey line.
